# An analysis of free energy methods to quantify and displace the KRasG12D Thr58 water

**DOI:** 10.1101/2025.04.12.648534

**Authors:** Callum J. Dickson

## Abstract

The KRasG12D mutant is an attractive target in oncology and can now be drugged via the switch-II allosteric pocket. Non-covalent ligands typically bind in the presence of a conserved structural water, which interacts with Thr58 and Gly10. In this work, we use a dataset of published non-covalent KRasG12D inhibitors to evaluate free energy methods for the interaction with or displacement of this conserved water molecule. We find that the Thr58 water and a second proximal water site are predicted to be unstable relative to bulk water and that relative binding free energy methods capture suitably well the binding affinities of ligands that disturb or replace these waters.

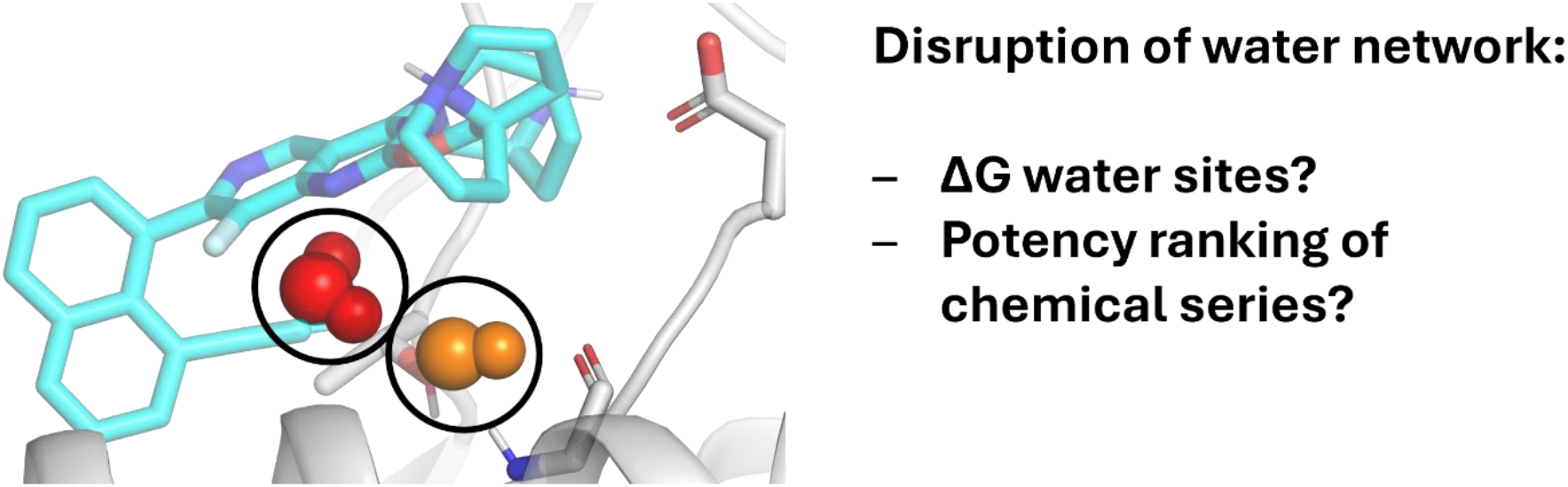

## Introduction

The RAS gene is one of the most frequently mutated gene families in cancers – the KRasG12C mutation is causally implicated in non-small cell lung cancer, whilst the KRasG12D mutation is most commonly occurring of the RAS mutations, arising in pancreatic ductal adenocarcinoma, colorectal cancer as well as non-small cell lung cancer.^1, 2^ Although previously considered undruggable, the discovery of the switch-II allosteric pocket by Shokat et. al.^3^ spurred the development of numerous small molecule covalent inhibitors of G12C which react with the cysteine-12 allele.^4-7^ More recently, small molecule inhibitors for the G12D mutant have also been reported, including inhibitors which non-covalently engage aspartic acid-12 with a basic amine.^8, 9^

MRTX1133 is one such G12D specific inhibitor, utilizing a bridged amine to engage aspartic acid-12.^8^ In the co-crystal structure of this compound bound non-covalently at the switch-II pocket of KRasG12D (PDB: 7RPZ), a conserved water molecule can be identified engaging in hydrogen bonding interactions to residues Gly10 and Thr58 of the protein while also receiving a CH hydrogen bond from the ethynyl group of the inhibitor. Indeed, this water has been observed to be present in the switch-II pocket of the majority of KRas structures in an analysis by Leini and Pantsar.^10^ Although selected KRasG12C covalent inhibitors displace this water productively, for example via the cyano group on MRTX849,^5^ reported G12D inhibitors to date are not able to displace this water in a manner that improves potency.

The understanding and displacement of “trapped” or energetically unfavourable water molecules has been of particular interest in the computational chemistry field. Numerous medicinal chemistry campaigns have successfully improved the potency of protein-ligand binding via the identification and displacement of energetically unfavourable water molecules proximal to ligand molecules undergoing lead optimization.^11-13^ As such, multiple algorithms have become available to identify water molecules in a ligand binding pocket and estimate their free energy, including WaterMap,^14, 15^ GIST,^16^ SSTMap,^17^ SZMAP,^18^ and WaterFLAP,^19^ among others.^20^ Furthermore, the importance of water sampling during relative binding free energy calculations has been highlighted.^21, 22^ Given that certain ligand transformations may result in groups occupying areas of the pocket containing trapped waters, the ability to sample movement of waters into bulk is important. Since trapped waters typically are unable to diffuse from the pocket during the timescales of molecular dynamics sampling, specific algorithms such as Grand Canonical Monte Carlo may be applied, which couples system waters to an external reservoir allowing insertion/deletion of water molecules.^21, 23^

In this work, we applied multiple methods to understand and quantify the energetics of the KRasG12D Thr58 water and selected proximal waters. We next used a dataset of published non-covalent KRasG12D inhibitors from the Mirati MRTX1133 patent (WO2021041671A1)^24^ which are identified to interact with or displace the Thr58 water in order to benchmark relative binding free energy methods and their ability to capture the energetics associated with the disruption of this conserved water molecule.

We find that although the Thr58 water is predicted to be displaceable by computational tools, to date non-covalent KRasG12D inhibitors have not been reported that efficiently displace this conserved water molecule. However, relative binding free energy methods are found to perform reasonably well within a dataset of molecules that disturb this water site, providing confidence that they could be utilized to identify suitable substituents able to displace the Thr58 in a productive manner.

## Methods

All calculations used the 7RT1 co-crystal structure of compound 15 bound to KRasG12D at resolution of 1.27 Å.^8^ Compound 15 is a close analog to patent example 186 (see Table 2), having a pyrrolidine side-chain in place of the butterfly amine and 2-naphthol. Both the Thr58 water (W1) and a second coordinating proximal water (W2) are evident in this structure (see Figure 1). This structure was prepared with CCG MOE protein preparation,^25^ assigning standard amino acid protonation states at pH 7.4, and optimization of the hydrogen bonding network. The 20 molecules extracted from WO2021041671A1^24^ (see Table 2) were prepared with Schrodinger ligprep and docked using GlideSP with maximum common substructure constraints to compound 15.^26, 27^

**Figure 1.**
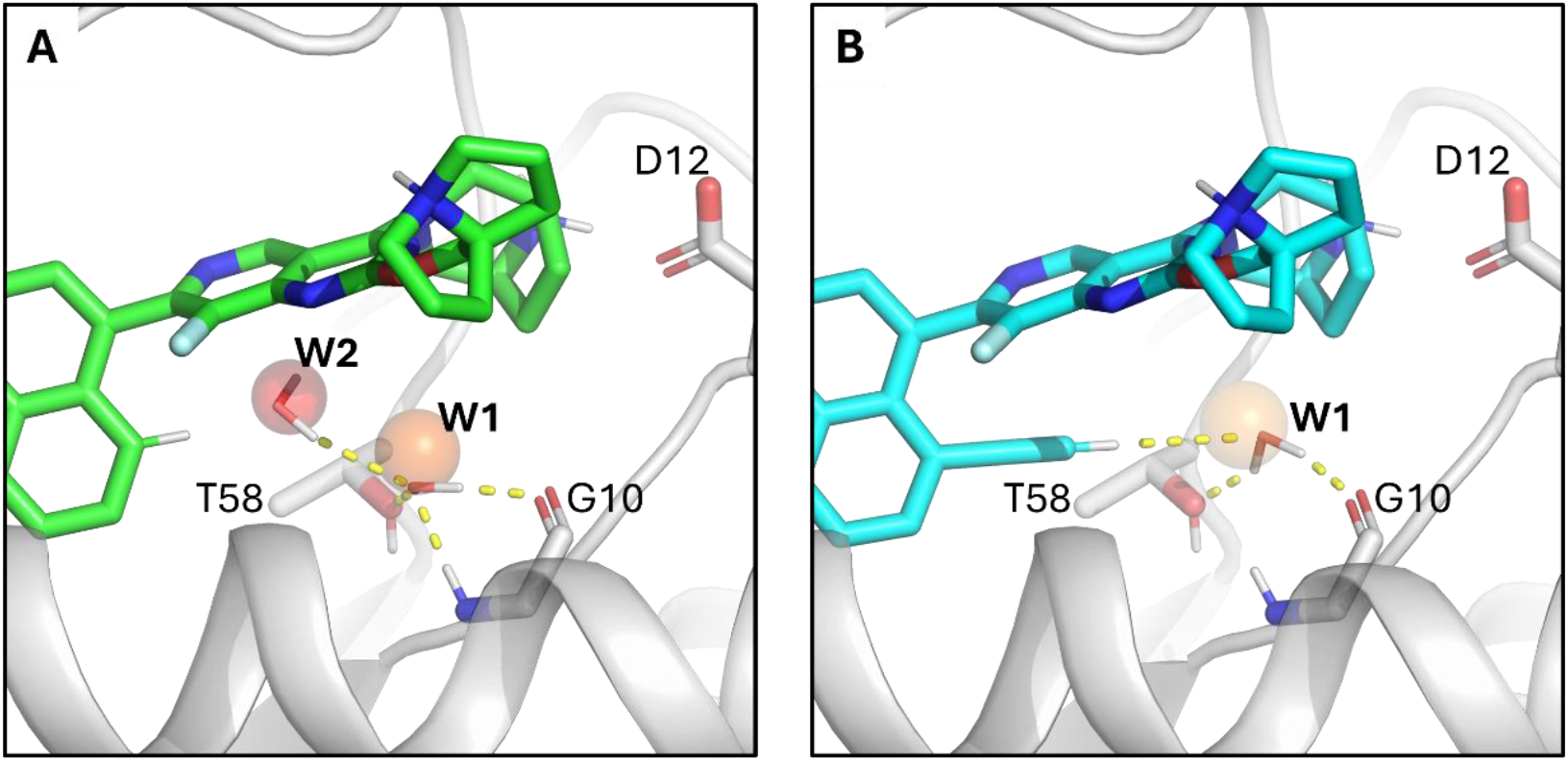
Binding models of A) ligand 186 and B) ligand 179 in the switch-II allosteric pocket of KRasG12D showing water sites W1 and W2 in the presence of ligand 186 and water site W1 only in the presence of ligand 179. The conserved water W1 receives a hydrogen bond from the backbone NH of Gly10 and is able to donate hydrogen bonds to Gly10 backbone carbonyl and Thr58 alcohol side-chain.

Water energetic calculations were performed using this system with either ligand 179 bound and only the Thr58 W1 water present, or ligand 186 bound and both W1 and W2 waters present (see Figure 1 and Table 2). WaterMap was performed using default settings of 2 ns sampling, the OPLS4 force-field^28^ and retaining the ligand, with waters within 10 Å of the ligand considered for analysis.^14, 15^ SSTMap calculations^16, 17^ were applied as a post-processing step to simulations run with Amber and PMEMD CUDA.^29-32^ After an equilibration of 3 ns NPT, systems were sampled for 10 ns with weak restraints of 2 kcal/mol/Å^2^ on backbone C-α atoms and 0.25 ps framerate. Proteins were modelled with ff19SB,^33^ ligands with gaff2^34^ and water molecules with TIP3P.^35^ All Amber MD simulations were performed in triplicate and results reported as bulk averages with standard deviation. Grand Canonical Monte Carlo (GCMC) simulations were performed with ProtoMS version 3.4.0.^23, 36^ The Amber ff14SB force-field was used to model proteins,^37^ ligands were modelled with gaff,^34^ and the water sphere containing the protein with the TIP4P force-field.^35^ Systems were equilibrated for 10 M steps before production sampling for 40 M steps at 298 K with a nonbonded cut-off of 10 Å. Systems were simulated in triplicate at 16 different *B* values equally spaced between -6.8 to -29.8 for ligand 186 and -0.8 to -23.8 for ligand 179 to control the chemical potential within a GCMC box of 96 Å^3^, containing the W1 and W2 sites.

All relative binding free energy calculations were performed in single edge star mode, using ligand 186 as the reference. FEP+ calculations were performed with default settings of 5 ns production sampling and 12 λ windows in the μVT ensemble using OPLS4 force-field.^28, 38^ AmberTI calculations were run in triplicate with a one-step protocol transforming both charges and van der Waals parameters between in the TI region of ligand 1 to ligand 2 with12 λ windows over a 5 ns production time as previously detailed.^39^ AmberTI runs used the ff14SB protein force-field,^37^ OpenFF 2.0 ligand force-field^40^ and OPC water model.^41^ The hydrogen mass repartitioning method^42^ was applied to allow a 4 fs timestep and the Metropolis Monte Carlo water sampling method used to sample water movements from trapped environments^43^ with nmc = 500 and nmd = 500. The timber package was used to prepare, run and analyse AmberTI simulations: https://github.com/callumjd/timber Results were analysed with python and scikit-learn to obtain performance metrics.^44^ Plots were created with matplotlib.^45^ Figures were generated with PyMOL.^46^

## Results & Discussion

### Energetics of Thr58 and proximal waters

Ligands in the MRTX1133 series bind the switch-II allosteric pocket at KRasG12D by engaging the aspartic acid-12 residue with a bridged amine, as seen in co-crystal structure 7RPZ.^8^ Within the pocket, two trapped waters W1 and W2 are evident in the des-ethyne analog compound 15 (7RT1) – see **Figure *1*** for docked pose of close analog ligand 186. Introduction of ethyne group at ligand 179 displaces water W2, leaving only the conserved Thr58 water W1. This water W1 receives a hydrogen bond from the backbone NH of Gly10 and is able to donate hydrogen bonds to Gly10 backbone carbonyl and Thr58 alcohol side-chain, in addition to receiving a weak CH hydrogen bond from the ethyne group of ligand 179.

We analysed the energetics of the Thr58 water (W1) at KRasG12D in the presence of ligand 179, having an ethyne group at the 8-position to form a CH hydrogen bond with W1 using SSTMap, WaterMap and ProtoMS (see Table 1). We also ran the same analyses in the presence of ligand 186, having a hydrogen at the 8-position, resulting in the presence of a second proximal water site W2. Given the difference in affinity between ligands 186 and 179 (ΔΔG^bind^ = -1.94 kcal/mol) the second water site W2 is expected to be energetically unfavourable. Although multiple ligands in the dataset (see Table 2) position substituents to displace both W1 and W2, the most potent inhibitor in the set is ligand 179, which displaces W2 and interacts directly with W1 via an ethyne group, suggesting that displacement of W1 from the MRTX1133 scaffold is difficult to achieve with the correct trajectory/functional group. Indeed, clinical candidate MRTX1133 retains the ethyne at the 8-position, differing from 179 only via an alcohol on the naphthyl and fluorine on the butterfly amine.^8^

**Table 1.** Water energy mapping results for sites W1 and W2 (occupancy, free energy and hydrogen bond count) for WaterMap, SSTMap and Grand Canonical Monte Carlo sampling for KRasG12D with either ligand 186 (hydrogen at naphthyl 8-position) or 179 (ethyne at naphthyl 8-position) bound in the switch-II allosteric pocket. All water site free energies are reported relative to bulk solvent.

|  | *186 bound* |  |  |  |  |  | *179 bound* |  |  |
| --- | --- | --- | --- | --- | --- | --- | --- | --- | --- |
| | W1 Occ | W1 $\Delta G$ (kcal/mol) | W1 HB | W2 Occ | W2 $\Delta G$ (kcal/mol) | W2 HB | W1 Occ | W1 $\Delta G$ (kcal/mol) | W1 HB |
| **WaterMap** | 0.99 | 5.34 | 2.59 | 0.99 | 9.23 | 1.25 | 0.98 | 9.11 | 1.96 |
| **SSTMap** | 0.86 ± 0.09 | 5.35 ± 0.16 | 3.15 ± 0.15 | 0.83 ± 0.09 | 5.5 ± 0.16 | 2.12 ± 0.37 | 0.91 ± 0.03 | 6.93 ± 0.21 | 2.21 ± 0.15 |
| **GCMC** | - | 3.43 ± 0.14 | - | - | 4.69 ± 0.18 | - | - | 5.14 ± 0.09 | - |

**Table 2.**
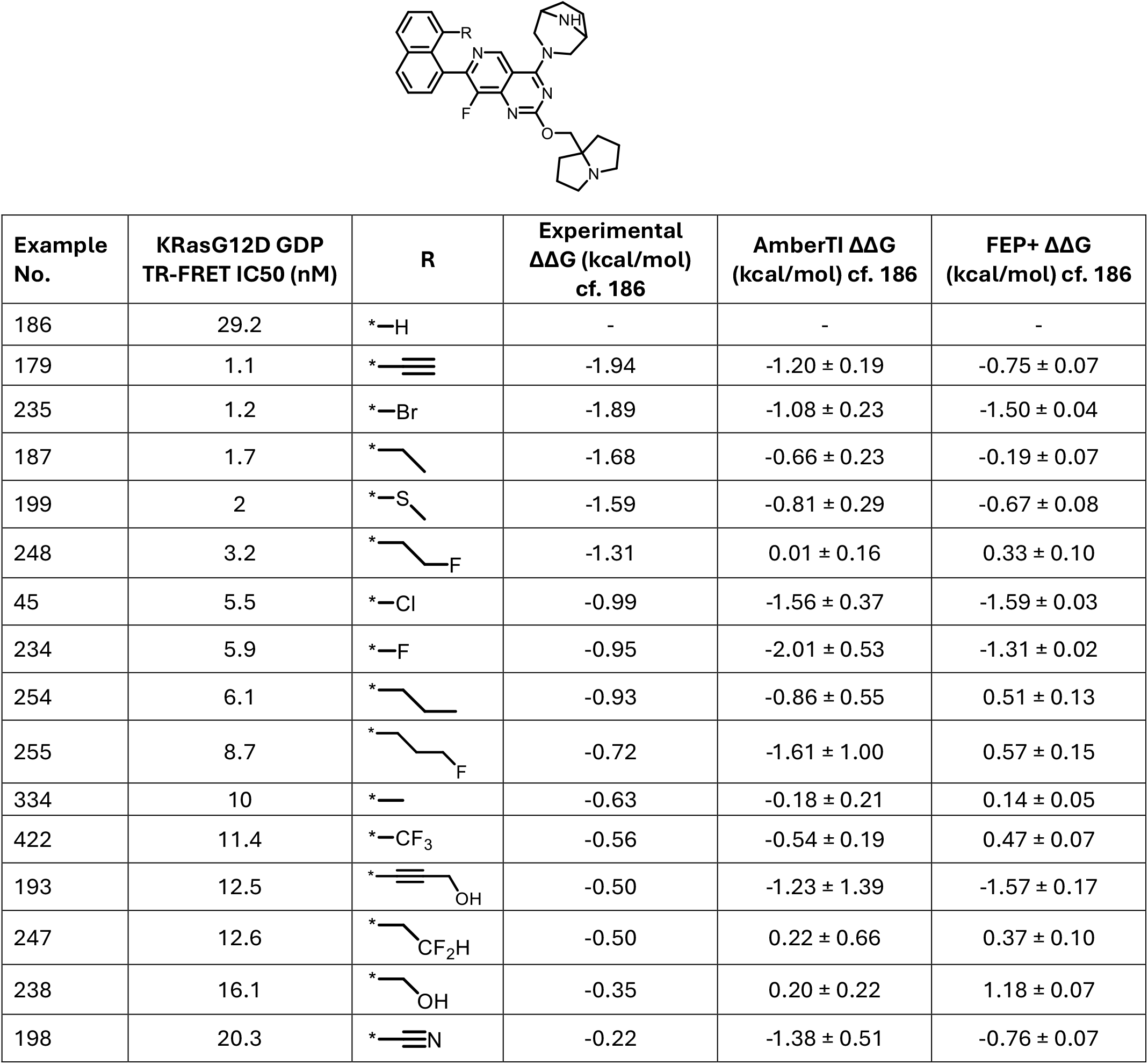
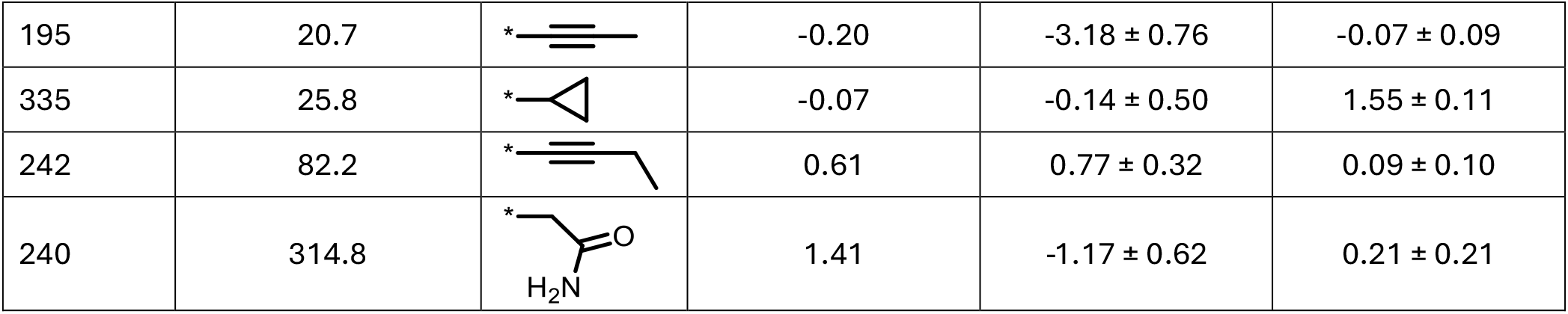
Dataset of KRasG12D inhibitors studied in this work with IC50 to inhibit KRasG12D in GDP loaded state from which the difference in binding free energy relative to Example 186 is reported.^24^ Also reported are the ΔΔGs relative to 186 calculated using AmberTI and FEP+ methods.

Analysis of the water mapping results (see Table 1) indicates that in all cases, the water mapping methods do rank water site W2 higher in free energy compared to W1 for simulations run in the presence of ligand 186. This is likely due to the lower number of hydrogen bonds water W2 is able to form. However, estimated free energies between the water sites vary, with WaterMap predicting a difference on the order of 4 kcal/mol between sites, while both SSTMap and GCMC find a lower difference between the W1 and W2 free energies.

In the case of simulations run in the presence of ligand 179, having an ethyne group to directly interact with W1, WaterMap, SSTMap and GCMC predict a destabilization of this water (the W1 ΔG is higher for 179 simulations compared to 186). This is likely due to the lower number of hydrogen bonds identified by the methods for water W1 in the presence of 179. The difference in free energy of binding between 186 and 179 is -1.94 kcal/mol, which may be attributed to the displacement of unstable water W2 and potentially stabilization of W1 via CH hydrogen bonds from the ethyne group to W1. Performing the following calculation:

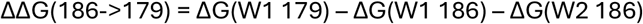

does predict a productive change in free energy of binding to displace water W2, despite the predicted increase in the free energy of water site W1, using WaterMap (−5.46 kcal/mol), SSTMap (−3.92 kcal/mol) and GCMC (−2.97 kcal/mol). The results are closest to the ΔΔG^bind^(186->179) = -1.94 kcal/mol for GCMC, however, this does not take into account the protein-ligand free energy change by introducing the ethyne. It should also be noted that the potential stabilization of W1 via the non-classical CH hydrogen bond from the ethyne of 179 may not be captured well by fixed charge molecular mechanics force-fields used for molecular dynamics and water mapping analysis.

The Grand Canonical Monte Carlo method is a rigorous method to understand the energetics of water networks in protein environments.^23^ The GCMC runs in the presence of 186 identify a water occupancy of 2 across W1 and W2 sites as expected, with one water site being 3.43 kcal/mol relative to bulk and the next site being less stable, 4.69 kcal/mol relative to bulk. The same simulations in the presence of ligand 179 predict an occupancy of 1 water only, again as expected, with W1 having a free energy prediction of 5.14 kcal/mol relative to bulk.

### Relative binding free energies of inhibitors interacting with or displacing Thr58 water

We studied a dataset of 20 ligands which extend different substituents at the 8-position of the naphthyl group towards the Thr58 residue (see Table 2).^24^ All relative binding free energy calculations were performed in star mode, transforming from ligand 186 as the reference, which has hydrogen at the 8-position.

We find reasonable performance of both AmberTI and FEP+ free energy methods (see Figure 2), with MUE<1 kcal/mol and RMSE<1.3 kcal/mol in both cases, the suggested RMSE range for prospective use.^47^ However, the correlation metrics for AmberTI are particularly poor, with r^2^=0.01 and Spearman ρ=0.17.

**Figure 2.**
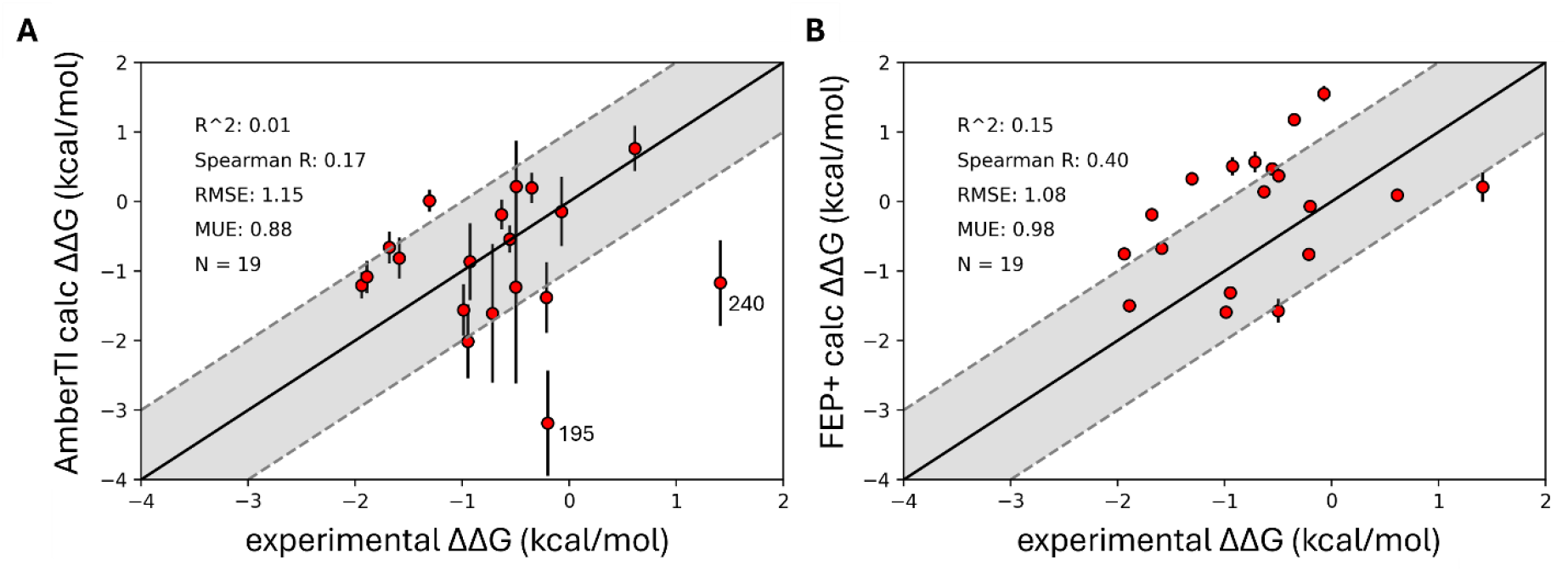
Performance of A) AmberTI and B) FEP+ relative binding free energy methods to predict the differences in affinity for KRasG12D inhibitors which extend substituents towards Thr58. In the case of AmberTI, two outliers are labelled.

An analysis of the outliers finds that in the case of AmberTI, ligand 195 which extends a butyne group towards Thr58 water and ligand 240, which extends a methyl amide into the Thr58 region are poorly predicted, with errors of >2.5 kcal/mol in both cases. In the case of ligand 195, interestingly both ligands 179 and 242 are well predicted by AmberTI, which differ only by one fewer and one additional methyl group, respectively. This may indicate that the subtleties of Thr58 water energetics are not captured in the 195 transformation. Although error bars are relatively large for ligand 195 (0.76 kcal/mol), the experimental value does not fall within the error range. In general, error bars on AmberTI ΔΔG predictions approach 1 kcal/mol for larger functional groups which occupy the Thr58 region such as 195 compared to smaller hydrophobic groups such as 45, 235 or 334.

In the case of FEP+, correlation metrics are improved over AmberTI, with r^2^=0.15 and Spearman ρ=0.40, although further improvements might be desired for prospective application to rank ordering. In general, transformations are under-optimistic from FEP+, with multiple transformations predicted above the +1 kcal/mol region beyond the identify line of Figure 2 B.

Overall, AmberTI and FEP+ relative binding free energy methods are in an acceptable range of RMSE <1.3 kcal/mol on this dataset of ligands which perturb the Thr58 region, suggesting they could be utilized to search for a substituent that is able to productively displace the Thr58 water.

## Conclusions

Displacement of trapped, energetically unfavourable water molecules is now a common strategy to improve the binding affinity of small molecule drugs in the hit-to-lead and lead optimization stages. In the case of KRasG12D, a conserved Thr58 water is present in solved co-crystal structures with MRTX1133 and related ligands. Despite efforts reported in the MRTX1133 patent, functional groups disturbing the Thr58 water do not improve affinity over the clinical candidate MRTX1133 which has an ethyne group forming a CH hydrogen bond to the Thr58 water molecule. In this work, we used water mapping methods to understand the energetics of this and proximal waters, finding that upon displacement of water site W2, the Thr58 W1 water is predicted to be further destabilized relative to bulk and potentially displaceable by an appropriate functional group. We next benchmarked two relative binding free energy methods, AmberTI and FEP+, for predictive performance against a dataset containing ligands that interact with or displace this water. Although rank correlation coefficients are poor, RMSE and MUE are in an acceptable range, suggesting such free energy of binding methods may be utilized in the search for appropriate chemical matter to displace the Thr58 water. In the case of AmberTI, substituents disturbing this water typically come with larger error between repeat calculations, with two ligands in particular giving rise to large outliers. Overall, computational methods capture the subtleties of Thr58 water energetics, giving confidence to utilize these methods in the search for substituents that are able to productively displace the Thr58 water in a manner that improves ligand binding affinity.

## Data availability

Input coordinates, FEP+ fmp files, SSTMap outputs, Amber topologies and an example TI lambda window are available for download here: https://github.com/callumjd/G12D_T58_water

